# Patterns of sedentary and active time accumulation are associated with mortality in US adults: The NHANES study

**DOI:** 10.1101/182337

**Authors:** Junrui Di, Andrew Leroux, Jacek Urbanek, Ravi Varadhan, Adam P. Spira, Jennifer Schrack, Vadim Zipunnikov

## Abstract

**Purpose:** Sedentary behavior has become a public health pandemic and has been associated with a variety of comorbidities including cardiovascular disease, type 2 diabetes, and some cancers. Previous studies have also shown that excessive amount of sedentary behavior is associated with all-cause mortality. However, no studies investigated whether patterns of sedentary and active time accumulation are associated with mortality independently of total sedentary and total active times. This study addresses this question by i) comparing several analytical ways to quantify patterns of both sedentary and active time accumulation through metrics of fragmentation of objectively-measured physical activity and ii) exploring the association of these metrics with all-cause mortality in a nationally representative US sample of elderly adults.

**Methods:** The accelerometry data of 3400 participants aged 50 to 84 in the National Health and Nutrition Examination Survey 2003-2006 cohorts were analyzed. Ten fragmentation metrics were calculated to quantify the duration of sedentary and active bouts: average bout duration, Gini index, average hazard, between-state transition probability, and the parameter of power law distribution. The association of these fragmentation metrics with all-cause mortality followed through December 31, 2011 was assessed with survey-weighted Cox proportional hazard models.

**Results:** In models adjusted for age, sex, race/ethnicity, education, body mass index, common comorbidities, and total sedentary/active time, four fragmentation metrics were associated with lower mortality risk: average active bout duration (HR=0.72 for 1SD increase, 95% CI = 0.59-0.88), Gini index for active bouts (HR = 0.75, 95% CI = 0.64-0.86), the parameter of power law distribution for sedentary bouts (HR = 0.75, 95% CI = 0.63-0.90), and sedentary-to-active transition probability (HR = 0.77, 95% CI = 0.61-0.96), and four fragmentation metrics were associated with higher mortality risk: the active-to-sedentary transition probability (HR = 1.40, 95% CI=1.23-1.58), the parameter of power law distribution for active bouts (HR = 1.33, 95% CI = 1.16-1.52), average hazard for durations of active bouts (HR = 1.32, 95% CI = 1.18-1.48), and average sedentary bout duration (HR =1.07, 95% CI = 1.01-1.13). After sensitivity analysis, average sedentary bout duration and sedentary-to-active transition probability became insignificant.

**Conclusion:** Longer average duration of active bouts, a lower probability of transitioning from active to sedentary behavior, and a higher normalized variability of active bout durations were strongly negatively associated with all-cause mortality independently of total active time. A larger proportion of longer sedentary bouts were positively associated with all-cause mortality independently of total sedentary time. The results also suggested a nonlinear association of average active bout duration with mortality that corresponded to the largest risk increase in subjects with average active bout duration less than 3 minutes.

## Introduction

Sedentary behavior is a significant risk factor for a wide range of chronic diseases, comorbidities, and mortality [1–10]. As people age, sedentary behaviors increase [7,11,12]. However, little is known about the detailed changes in daily sedentary patterns that accompany these shifts. A better understanding of temporal patterns of sedentary behavior in the characteristics of daily activities may clarify the role sedentary behaviors play in the progression of disability and disease, and thus provide more relevant public health recommendations regarding aging and diseases. Recently, the proliferation of wearable accelerometers has provided researchers with high-resolution, continuous activity data. As a result, accelerometer-measured physical activity (PA) offers the potential for both exploring detailed patterns of sedentary behaviors and providing more accurate estimates of overall sedentary behaviors, which have traditionally been underestimated by subjective methods [13,14].

Frequently, studies quantify sedentary behavior via an absolute amount (total sedentary minutes per day) or proportion (percentage) of waking hours spent sedentary [7,11,15,16]. Isotemporal substitution (ITS) model has been recently proposed to examine effects of replacing sedentary behavior with light PA (LiPA) and moderate-to-vigorous PA (MVPA) on weight, cardiovascular disease biomarkers, and mortality [2,17–19]. Compositional data analysis (CoDA) has been applied recently [20–22] to study the combined effects of time spent in LiPA and MVPA, sedentary behaviors and sleep while taking into account the codependence between those behaviors due to the finite time during a day. CoDA considers the time budget composition of the day without encountering issues of spurious correlations and collinearity. Both ITS and CoDA study the effect of allocation of the 24-hour budget between time spent in sedentary behavior, LiPA, MVPA and sleep and do not take into account how those total times have been accumulated. Several methods have been recently proposed to quantify the patterns of sedentary time accumulation to better understand how these patterns affect health and functional status. Conceptually, these methods segment objectively-measured daily activity into alternating bouts of sedentary and active time and the patterns are quantified via summaries of duration of frequency of switching between sedentary and active bouts.

From a statistical perspective, these methods can be grouped into two categories: nonparametric and parametric. Nonparametric approaches do not impose any distributional assumptions and summarize the distribution of bout durations via average duration, variability of durations, or describe properties of durations via hazard function. Paraschiv-Ionescu [23] used the area above the cumulative distribution function curve (AAC) of bout durations to study the relationship between chronic pain and PA. Mathematically, AAC is equivalent to the average bout duration. Healy et al. [5,24–26] and Chastin et al. [27] both proposed to use the reciprocal of average sitting or sedentary bout duration, which is shown in this paper to be related to the sedentary-to-active transition probability. The Gini index, a normalized measure of the variability of durations of sedentary bouts, has been proposed and applied by Chastin [13]. Lim [28] proposed to use two non-parametric summaries, *k_ar_* and *k_ra_*, to quantify the frequency of switching from active to resting (sedentary) and from resting (sedentary) to active state and defined them as the constant level of corresponding hazard functions.

Parametric approaches assume that bout durations follow a specific probability distribution, and then estimate the parameters that characterize the chosen distribution [13,23,27,29]. The most popular distribution used to quantify patterns of sedentary time accumulation is the power law distribution [13,23]. The parameter of power law has demonstrated stronger associations with clinical outcomes the total sedentary time in many clinical populations [13,23,27]. Another choice is the exponential distribution, a commonly used parametric distribution for time-to-event data [30]. Nakamura [29] modeled the duration of active periods using the stretched exponential distribution. Paraschiv-Ionescu [23] compared the performance of several heavy tail distributions to model sedentary and active duration, including the lognormal and the double Pareto distribution (Pareto2).

This work studies patterns of sedentary and active time accumulation with the fragmentation metrics outlined above using accelerometry data from 50 year and older participants of 2003-2006 National Health and Nutrition Examination Survey (NHANES). It compares the fragmentation metrics from both a practical and a statistical perspective and investigates whether these metrics are associated with mortality independently of total sedentary and total active time.

## Methods

### Study Populations and Measures

The National Health and Nutrition Examination Survey (NHANES) is a stratified, multistage, probabilistic sample representative of the civilian non-institutionalized U.S. population, described in detail elsewhere [31]. Fragmentation metrics were calculated from NHANES 2003-2004 and 2005-2006 waves, in which NHANES recruited a representative subsample aged 6 years and older to objectively evaluate PA using accelerometry. Since a survival analysis was conducted, subjects under the age of 50 years were excluded from the analysis to minimize potential biases induced by including younger individuals who were susceptible to genetic and/or atypical virulent diseases not related to habitual physical activity. NHANES data have been linked to death records from the National Death Index through December 31, 2011. If a participant is deceased, the duration of time in months between the NHANES examination and death is provided. Accidental deaths were excluded from the analysis. Demographic and comorbidity information that are available included, age, gender, race, education level, smoking status (never, former, current), drinking status (former or current drinker, heavy drinker, moderate drinker, non-drinker), body mass index (BMI; kg/m2), mobility difficulty (yes, no), diagnosis of diabetes, coronary heart disease, congestive heart failure, stroke, and cancer. Subjects with missing covariates were also excluded. In total, 3400 participants aged 50 to 84 years who fulfilled the inclusion criteria remained in the analysis, where 1773 were from cohorts 2003-2004, and 1627 were from cohorts 2005-2006. There were 542 reported deaths in the sample over an average of 6.4 follow-up years (7.2 for cohort 2003-2004, and 5.6 for cohort 2005-2006).

Physical activity was measured with the ActiGraph AM-7164 accelerometer (ActiGraph, LLC, Fort Walton Beach, Florida). The device was placed on an elasticized fabric belt, custom-fitted for each subject, and worn on the right hip. Participants were instructed to remove the belt while sleeping, bathing, and swimming. The monitors were programmed to record activity counts in successive 1-minute epochs for up to 7 consecutive days. Non-wear time was defined as any interval of 90 minutes or longer in which all count values were 0 with allowance for up to two minutes of non-zero counts between 1 and 99 [32,33]. Subjects were included if they had at least one valid day of accelerometer data, defined as at least 10 hours of wear time and all NHANES generated quality flags for the data were deemed valid [34].

Following previous work [35,36], minutes with activity counts < 100 are defined as sedentary, and minutes with activity counts ≥100 are defined as active without further distinguishing between light physical activity (LIPA) and moderate to vigorous physical activity (MVPA). A sedentary or active bout is defined as being sedentary or active for at least one minute.

### Notations

First, necessary notations are introduced. To define fragmentation metrics, the following notations are introduced: the duration of the longest active bout is denoted by D_*a*_, the number of bouts of length *t* is denoted by *n_A_(t)*, the number of active bouts of length *≤t* is denoted by *n_A_^c^(t)*. Total active time can be represented as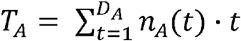, and the total number of active bouts can be represented as 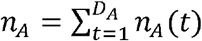. Notations for sedentary bouts can be defined similarly, with all subscripts changed to “S”. When introducing and defining the fragmentation metrics, the subscripts are sometimes dropped since they are defined for both sedentary and active. For simplicity of exposition, most of the conceptual derivations below will assume continuous *t*. Necessary adjustments are required to account for the fact that observed durations are integers. In this context, the continuity assumption is reasonable as a minute-level epoch assumption may be relaxed and a much finer resolution can be considered for high-frequency accelerometry data.

### Nonparametric metrics

We now review nonparametric fragmentation metrics.

#### 1. Average duration

The simplest and the most intuitive fragmentation metric is the average bout duration. Some previous studies have demonstrated usage of average bout durations. Paraschiv-Ionescu used the area above the cumulative distribution function curve (AAC) of bout durations (which statistically is equivalent to average duration) to study the relationship between chronic pain and PA [23]. To study how “breaks” of sedentary was associated with metabolic risk, Healy et al. considered the mean duration of the breaks [25]. Similarly, Lynch et al calculated the average length of active and sedentary bouts and studied their association with breast cancer biomarkers [37]. Average bout duration is denoted by *μ* and estimated as

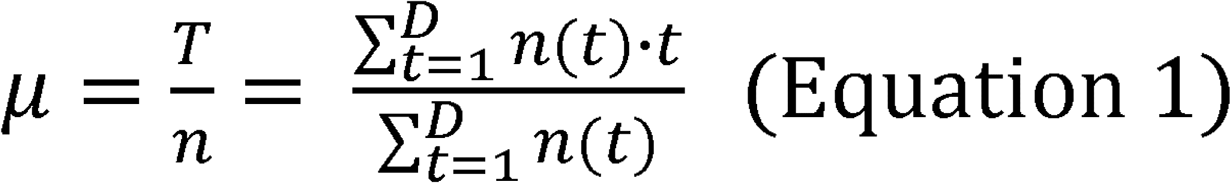

respectively.

#### 2. Normalized variability

Second most common method for nonparametrically summarizing a distribution is to estimate its variability. The Gini index was originally developed in econometrics to study the statistical dispersion of the distribution of incomes [38] and was used by Chastin et al [13] as a measure of the accumulation of sedentary time. Here, the Gini index is denoted by *g*, and defined and estimated as

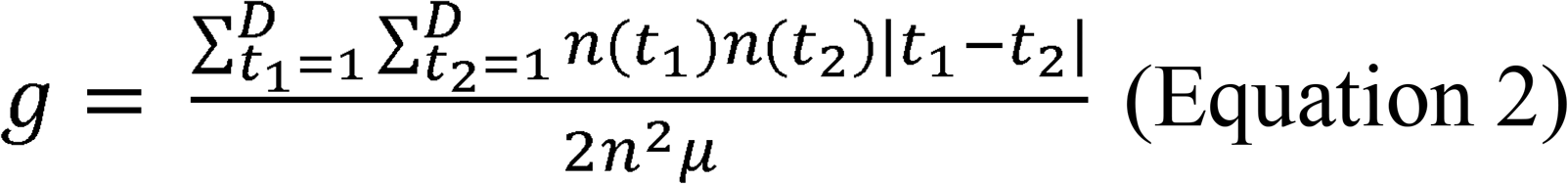

respectively.

The Gini index can be seen as a measure of (absolute, not squared) variability of bout durations normalized by the average duration. It can be shown that *g* is bounded between 0 and 1 [38]. When Gini index is close to 1, it indicates that total time is accumulated via a small number of longer bouts. Conversely, when Gini index is close to 0, it indicates that all bouts contribute equally to total time.

#### 3. Average hazard

Lim et al. [28] proposed two metrics *k_ra_* and *k_ar_* to quantify the frequency of switching from active to resting (sedentary) and from resting (sedentary) to active states, respectively, and studied their association with cognitive impairment in older adults [39]. Similarly, they adopted this concept to study sleep fragmentation and its effects on Alzheimer’s Disease (AD) [40,41]. The estimation procedure focused on estimation of “transition probability” as a function of bout duration, applied smoothing, and identified a range of durations when the “transition probability” function flattened out. The constant values of the “transition probability” from those “constancy” ranges have defined *k_ra_* and *k_ar_*.

From a statistical point of view, the “transition probability” function constructed by Lim et.al is exactly the hazard function widely used and studied in survival analysis. Modeling the hazard function is a principal approach for analyzing time-to-event data. The hazard function can be seen as the instantaneous probability of failure at time *t* given that the subject has survived until time *t* [30,42,43]. In this sense, the hazard is a measure of risk: the greater the hazard, the greater the risk of failure. Thus, in terms of dichotomous sedentary-active states, the hazard function can be used to study the probability of transitioning from sedentary to active or from active to sedentary state.

There are a few statistical reservations and modeling limitations to directly use the proposal of Lim et al. [28]. First and the most important, their proposal does not provide a well-defined estimand of interest. Second, the estimation of *k_ra_* and *k_ar_* tries to identify the “constancy range” of hazard function - a restrictive and hard-to-verify assumption. Third, the proposal employs LOWESS smoothing to identify the “constancy” range, thus, requires “ad hoc” decisions to make. To avoid these limitations, but stay close to the original proposal, we suggest the average hazard (AH) as the primary estimand to non-parametrically summarize the hazard function as a function of bout duration. Next, we outline our proposed procedure to estimate the AH.

For observed durations *t_1_,…,t_n_*, it is assumed that there are *m* unique values, which are denoted in increasing order by *{t_n1_,t_n2_*,…, *t_nm_}*. Then, hazard rates can be estimated at these distinct bout durations nonparametrically using the Nelson-Aalen approach [43] while treating all bout durations as non-censored (i.e, all active bouts will transition to sedentary bouts, and vice versa)

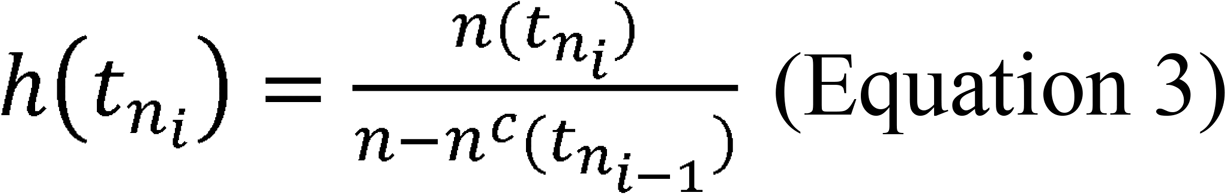

where *i = 1,…, m*. Note that Nelson-Aalen approach does not estimate the hazard function at time points that are not observed. The AH is then estimated as

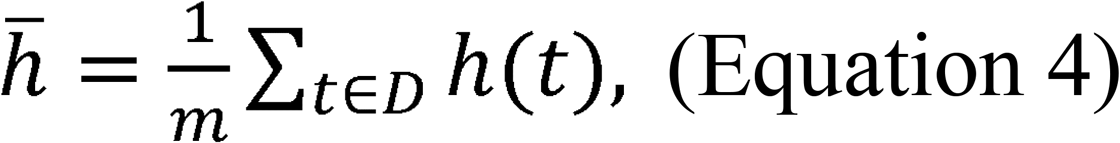

where D = *{t_n1_, t_n2_,…, t_nm_}*. Larger 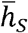 indicates a higher frequency of transitioning from sedentary to active state; and larger 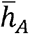 indicates a higher frequency of transitioning from active to sedentary state.

#### 4. Transition probability

Several studies have considered to use the reciprocal of average bout duration. To investigate the efficacy of a multicomponent intervention to reduce office workers’ sitting time, Healy et al. considered “sit to stand transition” which is defined as “number of sit-to stand transitions per hour of sitting” [24]. Chastin [27] studied the accumulation of sedentary time in older adults with obesity and low muscle strength by using the ratio of the number of sedentary bouts divided by the total sedentary time.

Here the reciprocal of average bout duration is denoted by *λ*. It can be shown (see proof in S1 Text) that *λ* is equal to between-states (i.e. sedentary-to-active or active-to-sedentary) transition

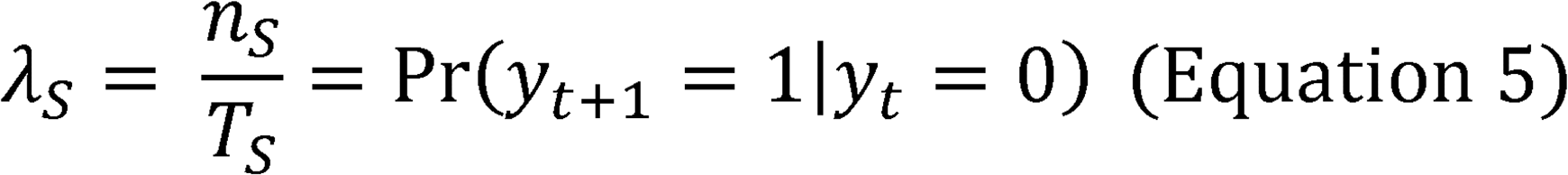

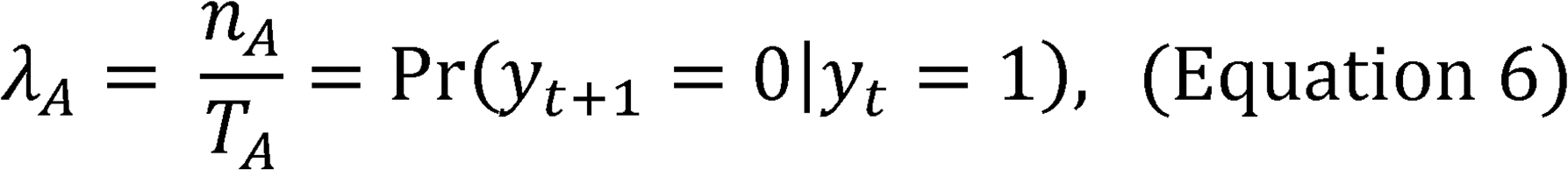

where *y_t_* is the indicator of activity type (sedentary or active) at time epoch *t*, and 0 and 1 denote sedentary and active states, respectively. Larger/smaller values of *λ* correspond to more/less frequent switching between the states and as a result, may indicate more/less fragmented activity pattern. It is important to note that larger/smaller values of *λ* also correspond to shorter/longer average bout duration.

From a parametric point of view, if the bout durations follow an exponential distribution, frequently used to model time-to-event data [30], *λ* is exactly the parameter that fully defines the exponential distribution with the following cumulative distribution function (CDF):

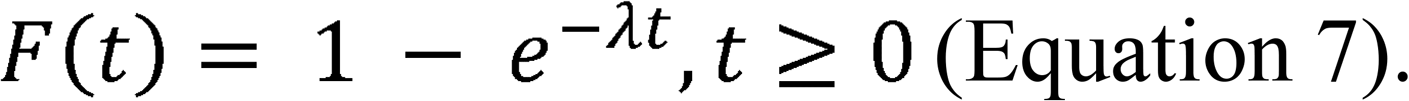

Note that any parametric assumptions should be validated through an appropriate goodness-of-fit tests.

### Parametric metrics

In this section, Power law, one of the most popular and widely used distribution to model bout duration, is discussed.

The CDF of a Power law or Type I Pareto distribution is defined as

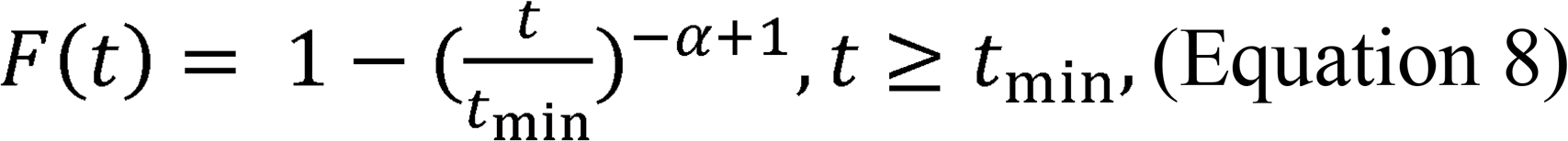

where *t_min_* is the lowest bout duration, the scaling parameter *α* summarizes information about the pattern of accumulation of total sedentary or total active time. Larger values of *α* indicates that subject tends to accumulate sedentary (active) time with a larger proportion of shorter sedentary (active) bouts.

The parameter *α* is typically estimated through maximum likelihood [44,45]. For the analysis of bout durations, the distribution is assumed to be discrete and the approximation to MLE estimator is used as follows

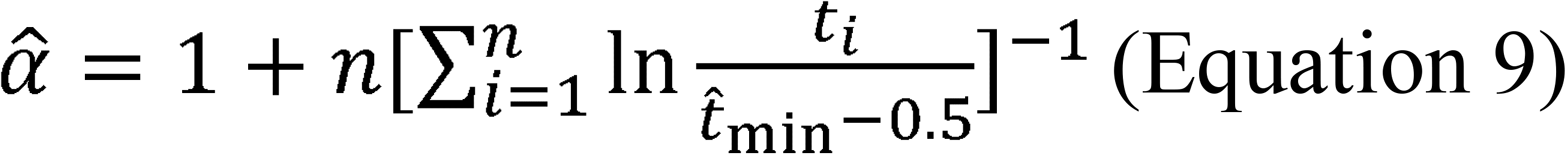

[44]. Here 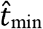 is the lower bound and can be estimated by the minimum bout length. Most studies [13,46] assess the goodness-of-fit of a power law through a visual assessment of the histogram of bout durations on doubly logarithmic plot. While Paraschiv-Ionescu employed a formal statistical goodness-of-fit test based on Anderson-Darling test [23].

Table 1 summarizes all five metrics described above along with their estimation method and interpretation.

**Table 1:**
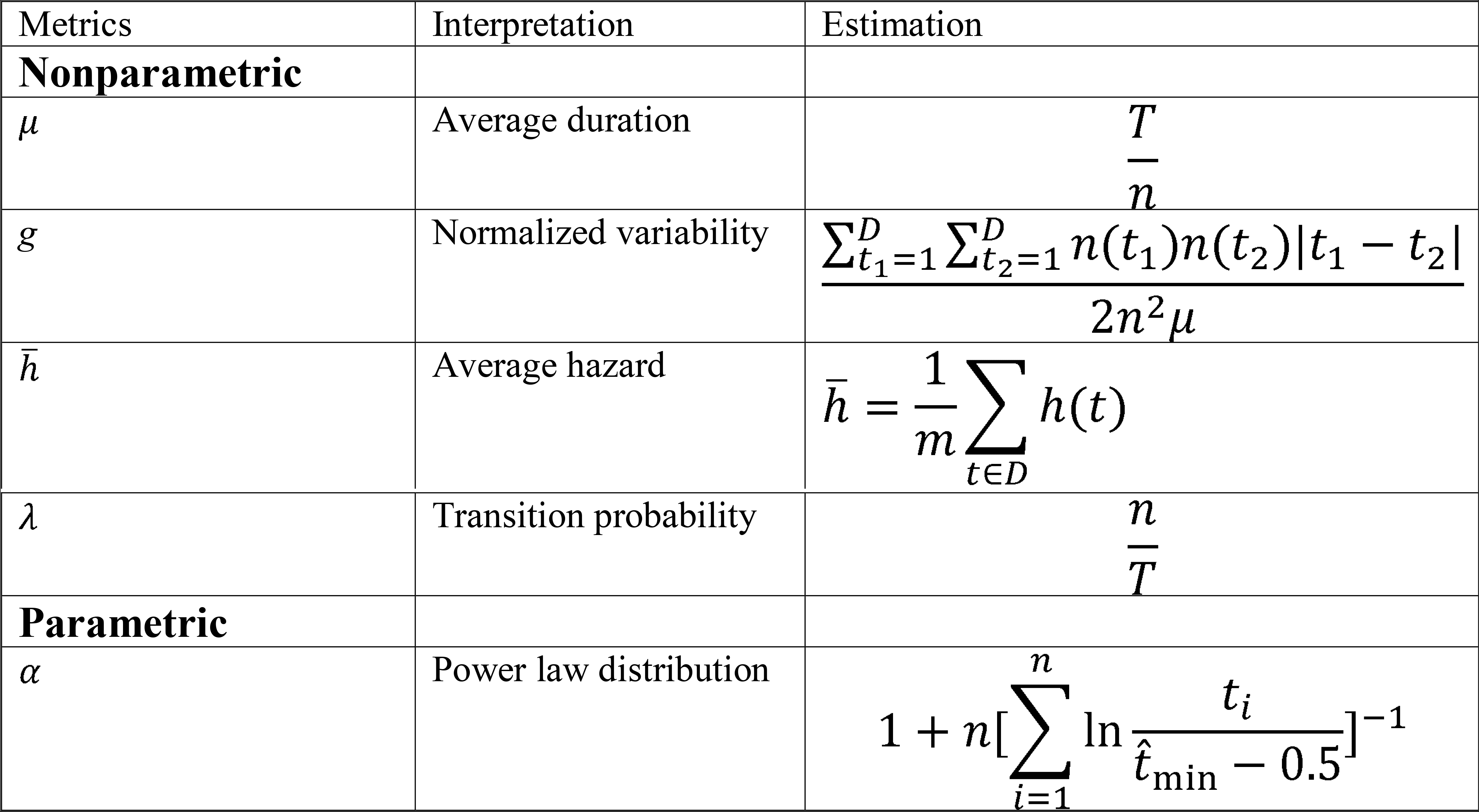
Summary of five-fragmentation metrics and their estimations

### Statistical analysis

Descriptive statistics were grouped by the survival status at the end of follow-up. For each subject, distributions of sedentary and active bouts were calculated by aggregating data from all bouts across valid days. Marginal densities for each metric was plotted by the survival status (deceased/alive). Survey-weighted Cox proportional hazard models [47–49] were fitted to model mortality. All models were adjusted for the covariates and comorbidities described in the Measures subsection. Two groups of models were fitted. Models in Group A studied the individual effect of each fragmentation metric on the relative risk of death by including one metric at a time. Models in Group B studied the individual effect of each metric on the relative risk of death independently of total sedentary/active times by including total sedentary time in the models with fragmentation metrics summarizing sedentary bouts and including total active time in the models with fragmentation metrics summarizing active bouts. For additional interpretability of the results, each fragmentation metric included in the models was standardized by subtracting population-level mean and dividing by population-level standard deviation, resulting in hazard ratios that correspond to one standard deviation change.

## Results

### Baseline characteristics

Descriptive statistics stratified by survival status at the end of follow-up period are presented in Table 2. All descriptive statistics are survey weighted to be representative of the U.S population. The average age of the participants was 64 years. Slightly more than half of participants were female (54%). Participants recorded as deceased tended to have greater daily sedentary time and lower daily active time. In addition, deceased participants tended to be older and have higher prevalence of comorbidities, mobility problems, and tobacco use.

**Table 2:**
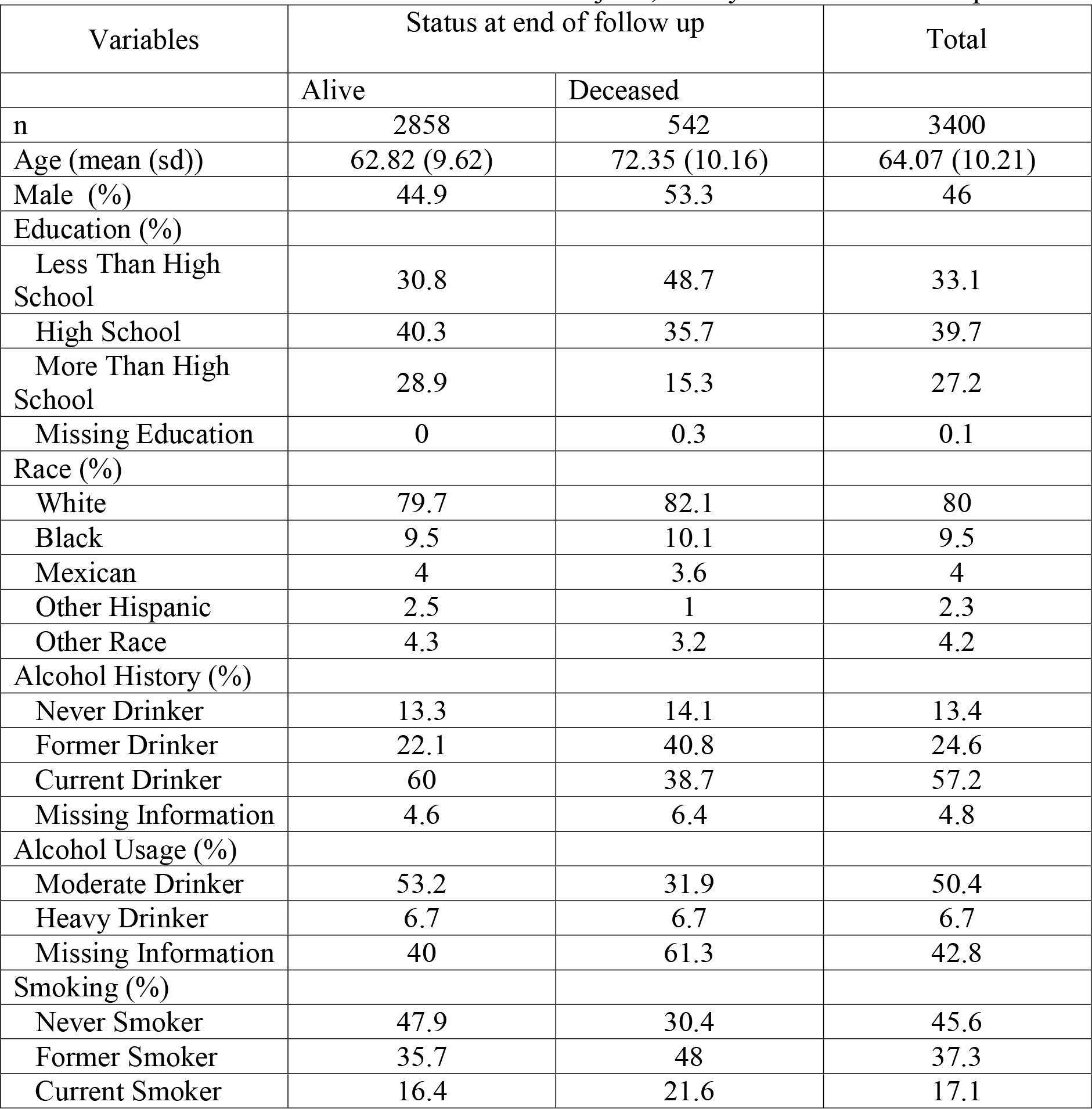
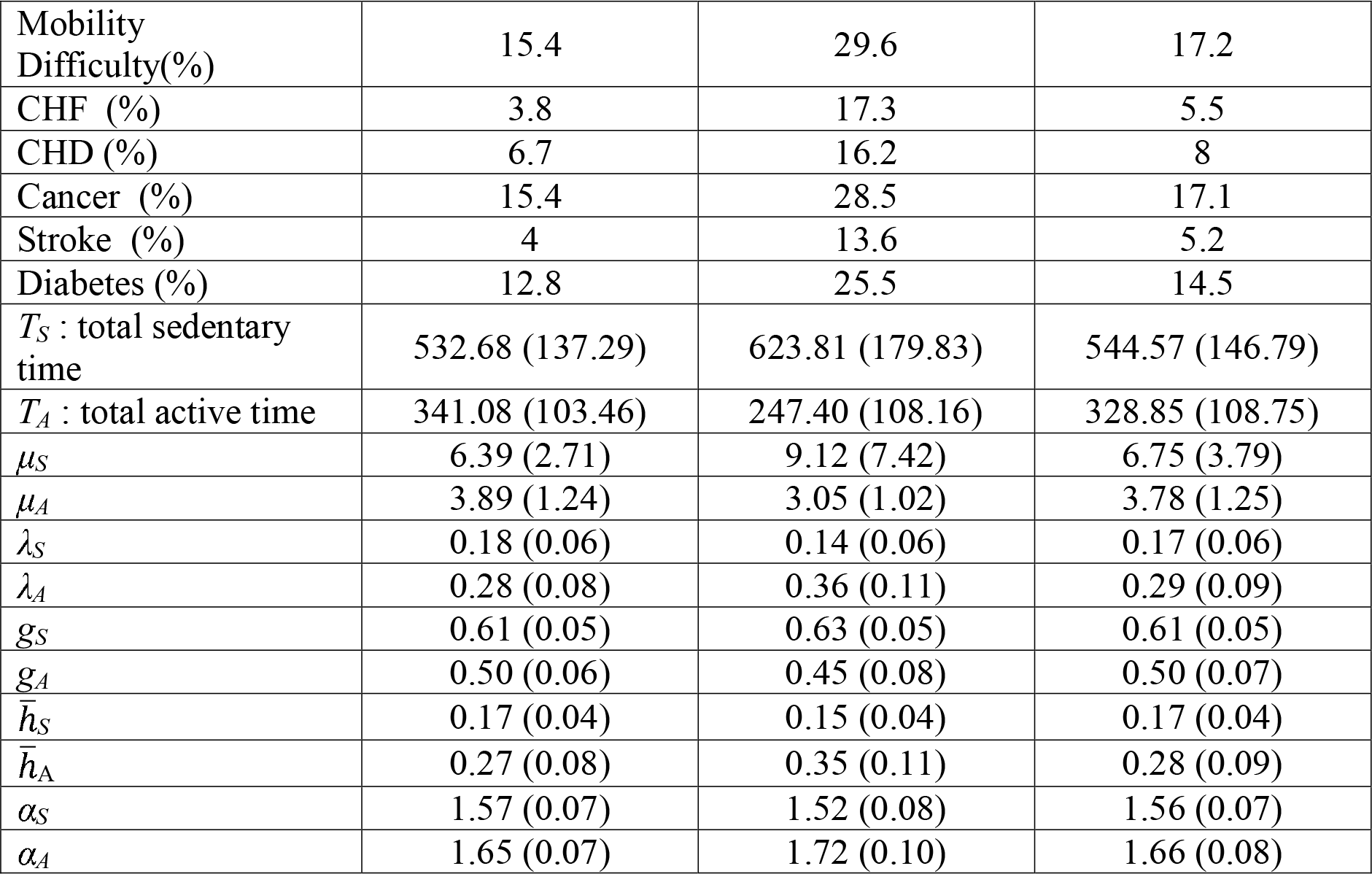
Baseline characteristics for all 3400 subjects, and by the end of follow-up status

| Variables | Status at end of follow up |  | Total |
| --- | --- | --- | --- |
|  | Alive | Deceased |  |
| n | 2858 | 542 | 3400 |
| Age (mean (sd)) | 62.82 (9.62) | 72.35 (10.16) | 64.07 (10.21) |
| Male (%) | 44.9 | 53.3 | 46 |
| Education (%) |  |  |  |
| Less Than High School | 30.8 | 48.7 | 33.1 |
| High School | 40.3 | 35.7 | 39.7 |
| More Than High School | 28.9 | 15.3 | 27.2 |
| Missing Education | 0 | 0.3 | 0.1 |
| Race (%) |  |  |  |
| White | 79.7 | 82.1 | 80 |
| Black | 9.5 | 10.1 | 9.5 |
| Mexican | 4 | 3.6 | 4 |
| Other Hispanic | 2.5 | 1 | 2.3 |
| Other Race | 4.3 | 3.2 | 4.2 |
| Alcohol History (%) |  |  |  |
| Never Drinker | 13.3 | 14.1 | 13.4 |
| Former Drinker | 22.1 | 40.8 | 24.6 |
| Current Drinker | 60 | 38.7 | 57.2 |
| Missing Information | 4.6 | 6.4 | 4.8 |
| Alcohol Usage (%) |  |  |  |
| Moderate Drinker | 53.2 | 31.9 | 50.4 |
| Heavy Drinker | 6.7 | 6.7 | 6.7 |
| Missing Information | 40 | 61.3 | 42.8 |
| Smoking (%) |  |  |  |
| Never Smoker | 47.9 | 30.4 | 45.6 |
| Former Smoker | 35.7 | 48 | 37.3 |
| Current Smoker | 16.4 | 21.6 | 17.1 |
| Mobility Difficulty(%) | 15.4 | 29.6 | 17.2 |
| CHF (%) | 3.8 | 17.3 | 5.5 |
| CHD (%) | 6.7 | 16.2 | 8 |
| Cancer (%) | 15.4 | 28.5 | 17.1 |
| Stroke (%) | 4 | 13.6 | 5.2 |
| Diabetes (%) | 12.8 | 25.5 | 14.5 |
| $T_S$ : total sedentary time | 532.68 (137.29) | 623.81 (179.83) | 544.57 (146.79) |
| $T_A$ : total active time | 341.08 (103.46) | 247.40 (108.16) | 328.85 (108.75) |
| $\mu_S$ | 6.39 (2.71) | 9.12 (7.42) | 6.75 (3.79) |
| $\mu_A$ | 3.89 (1.24) | 3.05 (1.02) | 3.78 (1.25) |
| $\lambda_S$ | 0.18 (0.06) | 0.14 (0.06) | 0.17 (0.06) |
| $\lambda_A$ | 0.28 (0.08) | 0.36 (0.11) | 0.29 (0.09) |
| $g_S$ | 0.61 (0.05) | 0.63 (0.05) | 0.61 (0.05) |
| $g_A$ | 0.50 (0.06) | 0.45 (0.08) | 0.50 (0.07) |
| $\bar{h}_S$ | 0.17 (0.04) | 0.15 (0.04) | 0.17 (0.04) |
| $\bar{h}_A$ | 0.27 (0.08) | 0.35 (0.11) | 0.28 (0.09) |
| $\alpha_S$ | 1.57 (0.07) | 1.52 (0.08) | 1.56 (0.07) |
| $\alpha_A$ | 1.65 (0.07) | 1.72 (0.10) | 1.66 (0.08) |

### Differences by survival status

Density plots of all metrics and total sedentary and active times categorized by the survival status are shown in Fig 1. These plots along with Table 2 estimates marginal associations between the fragmentation metrics and mortality status. Participants in the deceased group (red) tended to have longer sedentary time (T_*S*_) and shorter active time (T_*A*_), longer average sedentary bout *(μ_S_)* and shorter average active bout *(μ_A_)* durations than participants in the alive group (blue). Deceased participants were also more likely to have larger *g_S_* and smaller *g_A_*, indicating that their (normalized by the mean) sedentary bout durations are more variable while their (normalized by the mean) active bout durations are less variable. Considering the average hazard metrics, alive participants were more likely to transition from sedentary to active behavior (higher 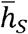), and deceased participants were more likely to transition from active to sedentary behavior (higher 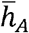). Deceased participants also had smaller *λ_S_* and higher *λ_A_* that corresponded to smaller chances of switching from sedentary to active behavior and higher chances of switching from active to sedentary behavior. Similarly, deceased participants tended to accumulate total active time with shorter active bouts (larger *α_A_*), and accumulate total sedentary time with longer sedentary bouts (smaller *α*s). Another interesting observation is that while the distributions of *μ_S_* and *μ_A_* exhibit considerable skewness, *λ_S_* and *λ_A_* had near-symmetric population-level distributions, a highly desirable statistical property. Note also that because of the skewness in distributions, the population averages of *μ_S_* and *μ_A_* are quite different from the reciprocals of the population averages of *λ_S_* and *λ_A_*.

**Fig 1.**
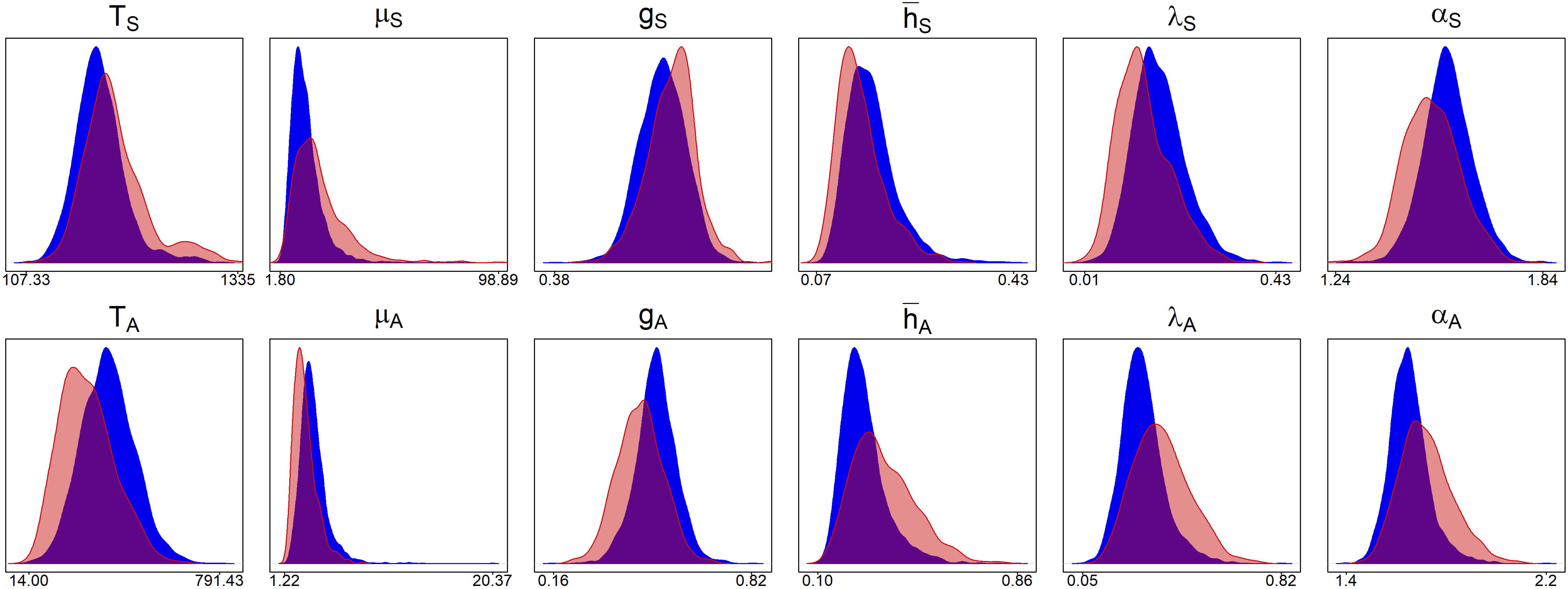
The estimated probability distribution functions of total sedentary time and sedentary fragmentation metrics (TOP) and total active time and active fragmentation metrics (BOTTOM) for deceased (red) and alive (blue) participants.

### Pairwise scatterplots and correlations of fragmentation metrics

Fig 2 shows the pairwise scatterplots (bottom triangle) and correlations (upper triangle) between all metrics. As expected, there was a clear parabolic association shape between μ and λ due to their definitions and estimation procedures. A parabolic relationship between μ and α was observed for both sedentary and active bouts. Meanwhile, λ and 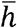 were highly positively correlated with a linear trend (*ρ = 0.84* for sedentary bouts, and *ρ = 0.93* for active bouts); and *λ* and *g* were highly negatively correlated with a linear trend (*ρ =−0.77* for sedentary bouts, and *ρ =−0.92* for active bouts). Moreover, *λ* and *α* had an almost linear relationship *ρ = 0.94* for sedentary bouts, and *ρ = 0.97* for active bouts). The correlation between the total sedentary time and the five sedentary bout fragmentation metrics were equal to 0.67, 0.57, −0.74, −0.74, −0.71. The correlations between the total active time and the five active bout fragmentation metrics were equal to 0.76, 0.74, −0.80, −0.81, −0.79. The shape and variability in the pair-wise scatterplots seem to indicate that fragmentation metrics may provide information about mortality beyond that of already provided by the total sedentary and total active times. In addition to revealing pairwise dependences between fragmentation metrics, Fig 2 allows to visually explore pairs of fragmentation metrics and their potential to separate deceased (red) from alive (blue) participants.

**Fig 2:**
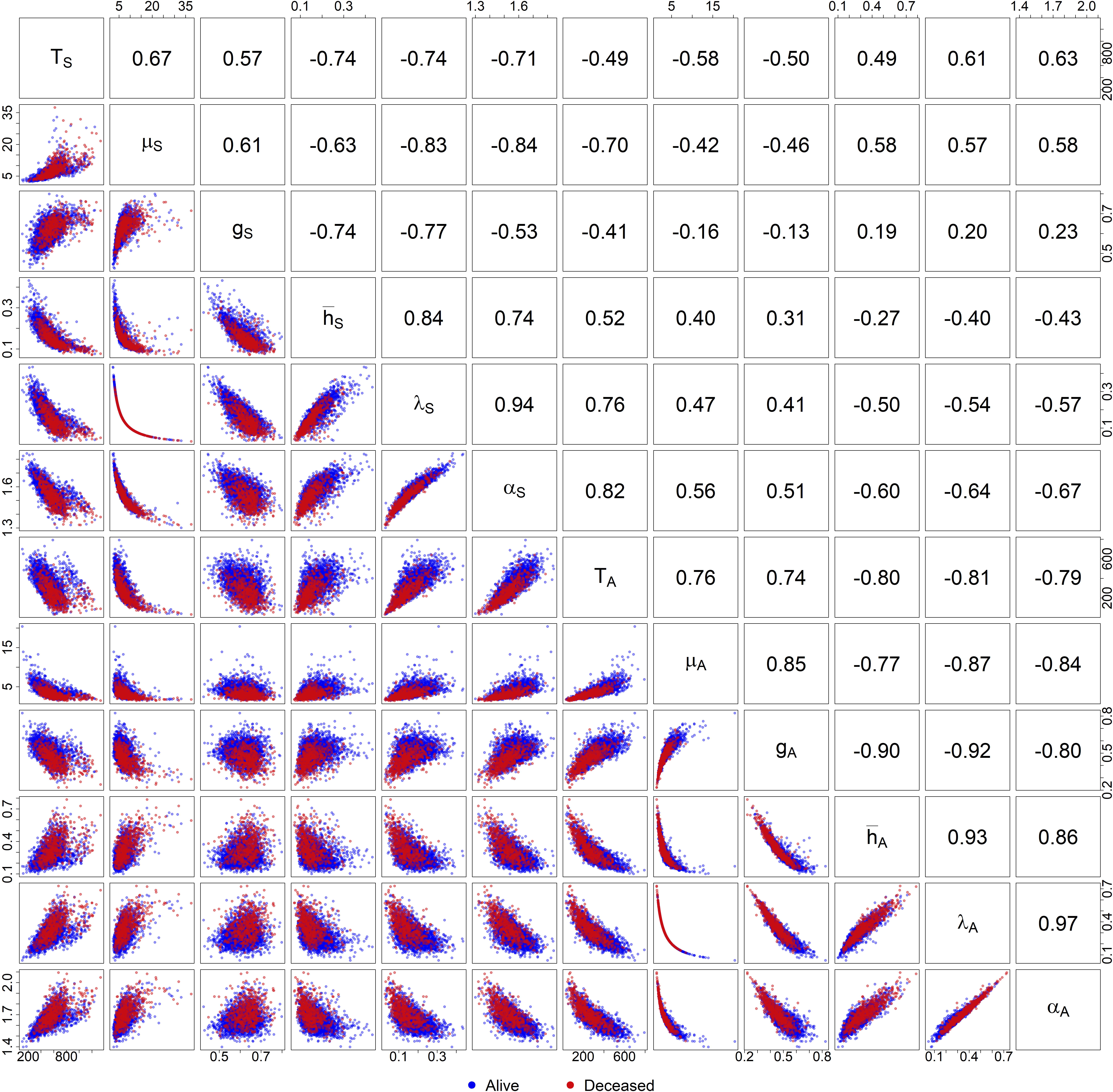
Pairwise scatterplots (lower triangular) and correlations (upper triangular) for the fragmentation metrics for deceased (red) and alive (blue). Deceased group was plotted over the alive group.

### Cox PH models

Table in S1 Table shows the results of the baseline models adjusted for all covariates and comorbidities described in Measures and adjusted for the total sedentary time (Model 1) or the total active time (Model 2). The results were consistent with those reported in previous studies [35,50,51] and demonstrate that one minute increase of total active time is associated with lower mortality risk (HR = 0.99, 95% CI = 0.99-1.00), and one minute increase of total sedentary time is associated with higher mortality risk (HR = 1.002, 95% CI = 1.001-1.002).

The left panel of Table 3 shows the odds ratio based on 1 SD increase for models in Group A (unadjusted for total sedentary and total active time). All ten fragmentation metrics were significantly associated with the relative odds of mortality. Five fragmentation metrics had a negative significant association with the relative odds of mortality: *μ*_A_(HR = 0.50, 95% CI = 0.42-0.60), *g_A_* (HR = 0.61, 95% CI = 0.55-0.69), 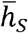 (HR = 0.80, 95% CI = 0.70-0.92), *λ_S_* (HR = 0. 67, 95% CI = 0.57-0.79), and *α_S_* (HR = 0.69, 95% CI = 0.60-0.78). Conversely, the other five fragmentation metrics had a positive association with the relative odds of mortality: *μ_S_* (HR = 1.11, 95 & CI = 1.05-1.16), *g_S_* (HR = 1.18, 95% 1.04-1.34), 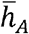 (HR = 1.52, 95% CI = 1.39-1.66), *λ_A_* (per SD HR = 1.59, 95% CI=1.45-1.75), and *α_A_* (HR = 1.56, 95% CI = 1.42-1.72).

**Table 3:**
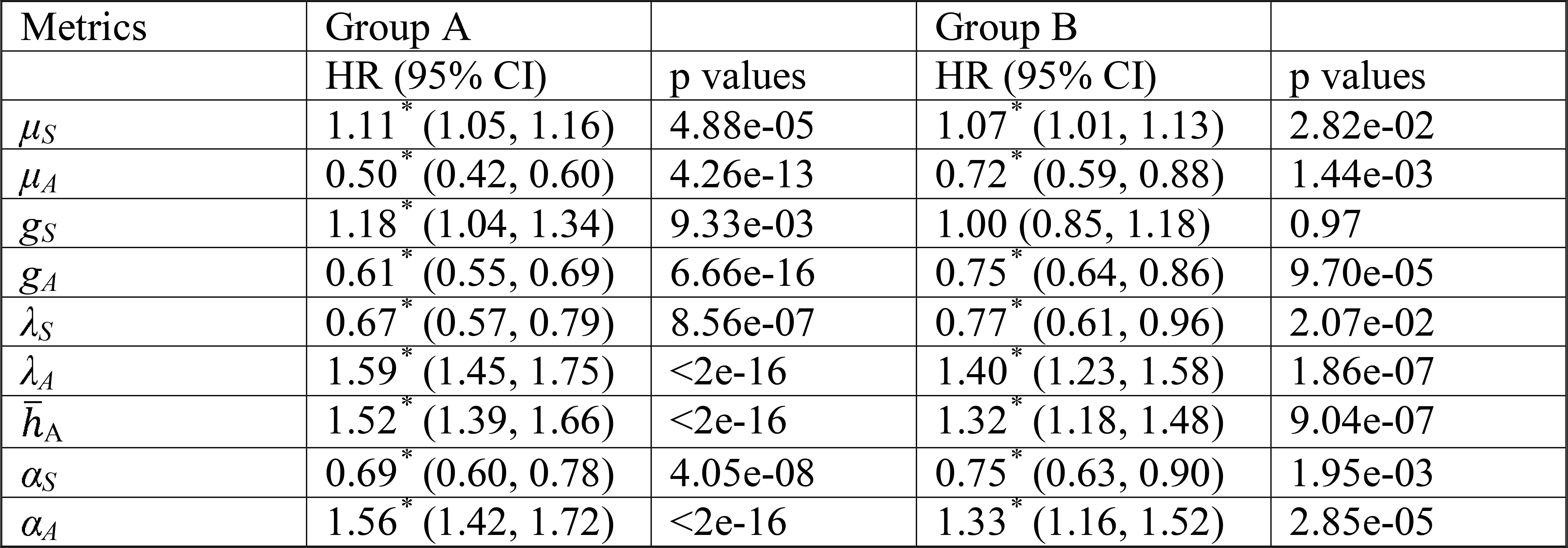
The results of fully adjusted Cox proportional hazard models (HR corresponding to 1SD increase)

| Metrics | Group A |  | Group B |  |
| --- | --- | --- | --- | --- |
|  | HR (95% CI) | p values | HR (95% CI) | p values |
| $\mu_S$ | 1.11* (1.05, 1.16) | 4.88e-05 | 1.07* (1.01, 1.13) | 2.82e-02 |
| $\mu_A$ | 0.50* (0.42, 0.60) | 4.26e-13 | 0.72* (0.59, 0.88) | 1.44e-03 |
| $g_S$ | 1.18* (1.04, 1.34) | 9.33e-03 | 1.00 (0.85, 1.18) | 0.97 |
| $g_A$ | 0.61* (0.55, 0.69) | 6.66e-16 | 0.75* (0.64, 0.86) | 9.70e-05 |
| $\lambda_S$ | 0.67* (0.57, 0.79) | 8.56e-07 | 0.77* (0.61, 0.96) | 2.07e-02 |
| $\lambda_A$ | 1.59* (1.45, 1.75) | <2e-16 | 1.40* (1.23, 1.58) | 1.86e-07 |
| $\bar{h}_A$ | 1.52* (1.39, 1.66) | <2e-16 | 1.32* (1.18, 1.48) | 9.04e-07 |
| $\alpha_S$ | 0.69* (0.60, 0.78) | 4.05e-08 | 0.75* (0.63, 0.90) | 1.95e-03 |
| $\alpha_A$ | 1.56* (1.42, 1.72) | <2e-16 | 1.33* (1.16, 1.52) | 2.85e-05 |

*Indicates significant association at 5% significance level.

HR: hazard ratio associated with one standard deviation increase

CI: confidence interval

The right panel of Table 3 shows the odds ratio based on 1 SD increase for models in Group B. Models in Group B additionally included total sedentary time in the models with sedentary bouts fragmentation metrics and included total active time in the models with active bouts fragmentation metrics. After the adjustment for the total sedentary/active times, four fragmentation metrics had a negative significant association with the relative odds of mortality: *μ_A_* (HR=0.72, 95 & CI = 0.59-0.88), *g_A_* (HR = 0.75, 95% CI = 0.64-0.86), *α_S_* (HR = 0.75, 95% CI = 0.63-0.90), *λ_S_* (per SD HR = 0.77, 95% CI = 0.62-0.96), and four fragmentation metrics had a positive significant association with the relative odds of mortality: *μ_S_* (HR =1.07, 95% CI = 1.011.13), 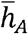 (HR = 1.32, 95% CI = 1.18-1.48), *α_A_* (HR = 1.33, 95% CI = 1.16-1.52), and *λ_a_* (HR = 1.40, 95% CI=1.23-1.58).

### Sensitivity Analysis

Two sensitivity analyses were conducted. The first explored possible effects of reversed causality by excluding all deaths within the first year of follow-up. The second excluded days with wear time longer than 20 hours to eliminate any bias from days when subjects wore accelerometers during sleep.

The first sensitivity analysis excluded deaths within the first year of follow-up. After the exclusion, 3334 (with 476 deaths) subjects remained and the same Cox PH models were reestimated. The results are shown in Table 4. The exclusion of the 1-st year follow-up deaths slightly attenuated results, but has not changed neither the significance or direction of most of the associations, with the exception of *λ_S_* that became insignificant in the model of Group B.

**Table 4:** The results of fully adjusted Cox proportional hazard models after excluding deaths within the first year of follow-up (HR corresponding to 1SD increase)

| Metrics | Group A |  | Group B |  |
| --- | --- | --- | --- | --- |
|  | HR (95% CI) | p values | HR (95% CI) | p values |
| $\mu_S$ | 1.11* (1.07, 1.16) | 7.18e-07 | 1.07* (1.02, 1.13) | 9.11e-03 |
| $\mu_A$ | 0.54* (0.45, 0.66) | 1.76e-09 | 0.71* (0.57, 0.87) | 1.30e-03 |
| $g_S$ | 1.15 (0.99, 1.34) | 0.056 | 0.98 (0.82, 1.17) | 0.83 |
| $g_A$ | 0.64* (0.57, .73) | 4.77e-13 | 0.74* (0.64, 0.86) | 1.01e-04 |
| $\bar{h}_S$ | 0.83* (0.73, 0.95) | 8.67e-03 | 1.05 (0.90, 1.24) | 0.52 |
| $\bar{h}_A$ | 1.45* (1.32, 1.59) | 1.48e-14 | 1.31* (1.16, 1.47) | 7.37e-06 |
| $\lambda_S$ | 0.73* (0.62, 0.85) | 4.92e-05 | 0.86 (0.69, 1.07) | 0.18 |
| $\lambda_A$ | 1.51* (1.37, 1.67) | <2e-16 | 1.39* (1.20, 1.60) | 5.75e-06 |
| $\alpha_S$ | 0.73* (0.64, 0.84) | 3.60e-06 | 0.82* (0.68, 0.99) | 0.04 |
| $\alpha_A$ | 1.48* (1.34, 1.63) | 7.66e-15 | 1.31* (1.13, 1.52) | 2.97e-04 |

The second sensitivity analysis excluded days with wear time longer than 20 hours. “Valid” days in the original analysis is defined based on the wearing time longer than 10 hour. However, there are numbers of subject-days that had more than 20 hours of wear time, with up to 24 hours of wear per day. Including days with wear time longer than 20 hours might lead to counting sleep time as sedentary that in turn could result in biased estimated summaries for sedentary and active time. Therefore, for the second sensitivity analysis, valid day was defined as a day with wear time between 10 and 20 hours. The threshold of 20 hours has been previously used to identify “bed-time” periods in NHANES [52]. After exclusion of invalid subject-days from the original samples, 3362 (with 534 deaths) subjects remained and same Cox PH models where re-estimated. The results are shown in Table 5. Although, the direction of association and significance for majority of the metrics remained the same, both *μ_S_* and *λ_S_* became insignificant.

**Table 5:** The results of fully adjusted Cox proportional hazard models after excluding invalid days with either too short or too long wear time (HR corresponding to 1SD increase)

| Metrics | Group A |  | Group B |  |
| --- | --- | --- | --- | --- |
|  | HR (95% CI) | p values | HR (95% CI) | p values |
| $\mu_S$ | 1.11* (1.05, 1.16) | 1.05e-04 | 1.05 (0.99, 1.11) | 0.13 |
| $\mu_A$ | 0.52* (0.43, 0.62) | 1.19e-11 | 0.78* (0.65, 0.95) | 1.24e-02 |
| $\lambda_S$ | 0.67* (0.58, 0.79) | 5.70e-07 | 0.79 (0.62, 1.02) | 0.07 |
| $\lambda_A$ | 1.54* (1.40, 1.70) | <2e-16 | 1.30* (1.14, 1.48) | 6.24e-05 |
| $g_S$ | 1.21* (1.06, 1.38) | 3.82e-03 | 1.05 (0.90, 1.22) | 0.53 |
| $g_A$ | 0.63* (0.56, 0.71) | 2.78e-14 | 0.78* (0.68, 0.90) | 5.23e-04 |
| $\bar{h}_S$ | 0.84* (0.74, 0.96) | 8.64e-03 | 1.03 (0.91, 1.16) | 0.68 |
| $\bar{h}_A$ | 1.49* (1.36, 1.63) | <2e-16 | 1.27* (1.15, 1.41) | 2.03e-06 |
| $\alpha_S$ | 0.70* (0.61, 0.80) | 3.15e-07 | 0.79* (0.63, 1.00) | 4.99e-02 |
| $\alpha_A$ | 1.51* (1.37, 1.66) | <2e-16 | 1.23* (1.08, 1.42) | 2.62e-03 |

### Nonlinear effect of average active bout and average sedentary bout durations

Among all considered fragmentation metrics, average bout duration, *μ*, is probably the most straightforward to calculate and communicate. Below, we show that, when modeled via *λ (=1/μ)*, the effect of *μ* on proportional HR is highly nonlinear and the largest risk increase is observed in subjects with average active bout duration less than 3 minutes.

To illustrate our argument, we refer to Table in S2 Table that reports estimated effect for non-standardized fragmentation metrics with respective to 1 unit increase for *μ_A_* and *λ_A_*, which equal to −0.26 and 3.88, respectively. The top panel in Fig 3 shows the hazard ratios with respect to a 1 minute increase of average activity bout calculated based on *μ_A_*, (dashed line) and *λ_A_* (solid line). When estimated from the Cox-PH model with *μ_A_*, the effect on the hazard ratio is constant and equal to e^−027^ = 0.77 (dashed line in the top panel). This implies that increasing average active bout by 1 minute always reduces hazard by approximately 23%, regardless of “original” average bout duration. On contrary, when considering the effect of increasing *μ_A_* by 1 minute estimated in the Cox-PH model with *λ_A_*, the nonlinear change is given by

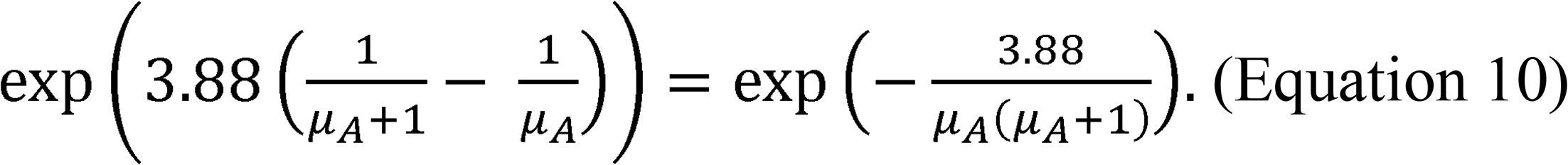

**Fig 3.**
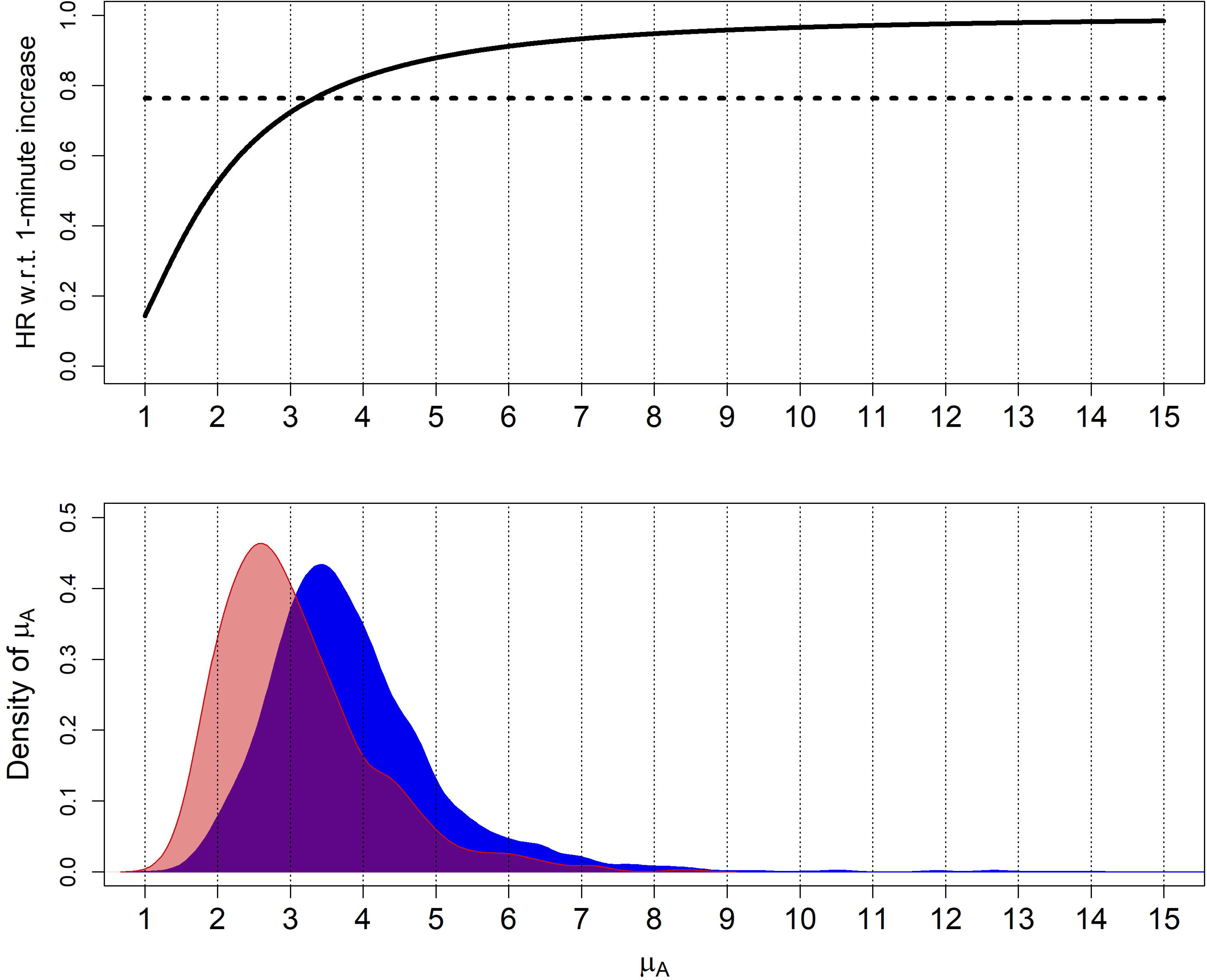
Effect of the change in *μ_A_* on hazard ratio. Top: Change in hazard ratios as a function of 1-minute change in *μ_A_*, based on *μ_A_* (dashed) and *λ_A_* (solid). Bottom: density of *μ_A_* for deceased (red) and alive (blue).

The solid black line on top panel of Fig 3 demonstrates this nonlinear effect on the hazard ratio. This shows that effect is drastically large and nonlinear when *μ_A_* is small, and as *μ_A_* gets larger, the effect flattens out. For example, if *μ_A_* = 1, then the hazard ratio corresponding to one-minute-increase is 0.14, i.e. increasing from a 1-minute to 2-minute average active bout makes hazard almost 86% (or 7 times) smaller. However, if *μ_A_* = 4, a one-minute increase in the average active bout reduces hazard ratio to 0.82, which translates to roughly 18% decrease. The observed non-liner effect is consistent with the density plots of average active bout durations shown at the bottom panel of Fig 3. It also shows that participants with average active bout shorter than 3 minutes are at highest risk of non-surviving.

## Discussion

To our knowledge, this study is the first that demonstrated that patterns of sedentary and active time accumulation are strongly associated with all-cause mortality independently of total sedentary and total active time, respectively. The major findings suggest that patterns of active time accumulation have much stronger significant associations with mortality than patterns of sedentary time accumulation. Longer average duration of active bouts, a lower probability of transitioning from active to sedentary behavior, and a higher normalized variability of active bout durations were strongly negatively associated with all-cause mortality independently of total active time. Although, the main analysis showed that a longer average duration of sedentary bouts and a lower probability of transitioning from sedentary to active behavior were positively associated with all-cause mortality, these associations became non-significant during sensitivity analysis. A larger proportion of longer sedentary bouts remained positively significantly associated with all-cause mortality during both sensitivity analyses.

The results for durations of active bouts suggest nonlinear associations of average active bout duration with mortality. The largest nonlinear risk increase was observed in subjects with average active bout duration less than 3. These results are consistent with previous studies showing positive health effects of accumulating active time in prolonged active bouts [25,37], but provide more insight in developing further individualized guidelines of physical activity.

The results regarding sedentary bout duration may imply that subjects that accumulate their sedentary time through prolong sedentary bouts may benefit from breaking those long bouts.

This is consistent with the emerging evidence that breaking up prolong sedentary time can have multiples positive effects including improvement of cardiovascular and cardiometabolic health [3,5,6,24,26,53,54].

Methodologically, this work compared various approaches to quantify patterns of sedentary time accumulation through fragmentation of accelerometry measured physical activity. Active-to-Sedentary and Sedentary-to-Active transition probabilities, co-expressed as reciprocals of average active bout and average sedentary bout durations, are easily interpretable modifiable fragmentation metrics with insightful connection to other metrics of fragmentation, and stronger associations to mortality. Importantly, the use of *λ* results in a non-linear effect on hazard ratio that could potentially help with providing more subject-specific guidelines. The parameter of power law distribution, α, has been a popular metric in a few recent studies on accumulation patterns of sedentary time [13,46,53]. Fig 2 demonstrates a close-to-linear relationship between *α* and *λ* with correlation coefficients of 0.94 for sedentary bouts and 0.97 for active bouts. Nevertheless, *α_S_* was the only sedentary time fragmentation metric that remained significant through the sensitivity analyses. This supports recent findings where among several sedentary fragmentation metrics only *α_S_* was significantly associated with biomarkers of glucose metabolism [53].

There are a few limitations in the present study. First, the minimum bout length was defined based on the Actigraph epoch duration of 1 minute. The results may change, if the sedentary and active bouts are defined in a different way. Second, the threshold of 100 AC used to define sedentary state is a widely accepted for the NHANES data. However, there is an on-going debate on whether this threshold can be uniformly applied across all age groups and genders and whether such uniform thresholds induce bias in classifying sedentary behaviors. Therefore, it is imperative to fragmentation metrics as a function of different threshold values in future work. Third, the non-wear criteria may overestimate the total amount of sedentary time by including those have not complied with the protocol and wear the device during sleep time. Finally, this study is cross-sectional, therefore, all findings are associative, not casual.

Fragmentation metrics can provide unique translatable insights into accumulation patterns for sedentary and active time and lead to the better understanding of associations between those patterns and health outcomes.

## Acknowledgements

## Supporting Information

**S1 Table. Results of the baseline models adjusted for all covariates and comorbidities described in Measures and adjusted for the total sedentary time (Model 1) or the total active time (Model 2).**

**S2 Table. Results of Cox PH models in original scale without standardization.**

**S1 Text. Heuristic Proof that is equivalent to the between states transition probability.**

**S2 Text. Expressing nonparametric metrics using the empirical CDF.**

## References

1. Varo JJ, Martínez-González MA, de Irala-Estévez J, Kearney J, Gibney M, Martínez JA. Distribution and determinants of sedentary lifestyles in the European Union. Int J Epidemiol. 2003;32: 138–146. doi: 10.1093/ije/dyg116

2. Buman MP, Winkler EAH, Kurka JM, Hekler EB, Baldwin CM, Owen N, et al. Reallocating Time to Sleep, Sedentary Behaviors, or Active Behaviors: Associations With Cardiovascular Disease Risk Biomarkers, NHANES 2005-2006. Am J Epidemiol. 2014;179: 323–334. doi:10.1093/aje/kwt292

3. Owen N, Healy GN, Matthews CE, Dunstan DW. Too much sitting: the population-health science of sedentary behavior. Exerc Sport Sci Rev. 2010;38: 105–113. doi:10.1097/JES.0b013e3181e373a2

4. Hamilton MT, Healy GN, Dunstan DW, Zderic TW, Owen N. Too little exercise and too much sitting: Inactivity physiology and the need for new recommendations on sedentary behavior. Curr Cardiovasc Risk Rep. 2008;2: 292–298. doi: 10.1007/s12170-008-0054-8

5. Dunstan DW, Kingwell BA, Larsen R, Healy GN, Cerin E, Hamilton MT, et al. Breaking Up Prolonged Sitting Reduces Postprandial Glucose and Insulin Responses. Diabetes Care. 2012;35: 976–983. doi:10.2337/dc11-1931

6. Owen N, Sparling PB, Healy GN, Dunstan DW, Matthews CE. Sedentary Behavior: Emerging Evidence for a New Health Risk. Mayo Clin Proc. 2010;85: 1138–1141. doi:10.4065/mcp.2010.0444

7. Matthews CE, Chen KY, Freedson PS, Buchowski MS, Beech BM, Pate RR, et al. Amount of Time Spent in Sedentary Behaviors in the United States, 2003-2004. Am J Epidemiol. 2008;167: 875–881. doi:10.1093/aje/kwm390

8. Chau JY, Grunseit AC, Chey T, Stamatakis E, Brown WJ, Matthews CE, et al. Daily Sitting Time and All-Cause Mortality: A Meta-Analysis. Gorlova OY, editor. PLoS One. 2013;8: e80000. doi:10.1371/journal.pone.0080000

9. Matthews CE, George SM, Moore SC, Bowles HR, Blair A, Park Y, et al. Amount of time spent in sedentary behaviors and cause-specific mortality in US adults. Am J Clin Nutr. 2012;95: 437–445. doi:10.3945/ajcn.111.019620

10. Healy GN, Clark BK, Winkler EAH, Gardiner PA, Brown WJ, Matthews CE. Measurement of Adults’ Sedentary Time in Population-Based Studies. Am J Prev Med. 2011;41: 216–227. doi:10.1016/j.amepre.2011.05.005

11. Davis MG, Fox KR, Hillsdon M, Sharp DJ, Coulson JC, Thompson JL. Objectively Measured Physical Activity in a Diverse Sample of Older Urban UK Adults. Med Sci Sport Exerc. 2011;43: 647–654. doi:10.1249/MSS.0b013e3181f36196

12. Varma VR, Dey D, Leroux A, Di J, Urbanek J, Xiao L, et al. Re-evaluating the effect of age on physical activity over the lifespan. Prev Med (Baltim). 2017; 101: 102–108. doi:10.1016/j.ypmed.2017.05.030

13. Chastin SFM, Granat MH. Methods for objective measure, quantification and analysis of sedentary behaviour and inactivity. Gait Posture. 2010;31: 82–86. doi:10.1016/j.gaitpost.2009.09.002

14. Reilly JJ, Penpraze V, Hislop J, Davies G, Grant S, Paton JY. Objective measurement of physical activity and sedentary behaviour: review with new data. Arch Dis Child. 2008;93: 614–619. doi:10.1136/adc.2007.133272

15. Hamilton MT, Hamilton DG, Zderic TW. Role of Low Energy Expenditure and Sitting in Obesity, Metabolic Syndrome, Type 2 Diabetes, and Cardiovascular Disease. Diabetes. 2007;56: 2655–2667. doi:10.2337/db07-0882

16. Zderic TW. Physical inactivity amplifies the sensitivity of skeletal muscle to the lipid-induced downregulation of lipoprotein lipase activity. J Appl Physiol. 2006;100: 249–257. doi: 10.1152/japplphysiol.00925.2005

17. Schmid D, Ricci C, Baumeister SE, Leitzmann MF. Replacing Sedentary Time with Physical Activity in Relation to Mortality. Med Sci Sport Exerc. 2016;48: 1312–1319. doi:10.1249/MSS.0000000000000913

18. Rossen J, Buman MP, Johansson U-B, Yngve A, Ainsworth B, Brismar K, et al. Reallocating bouted sedentary time to non-bouted sedentary time, light activity and moderate-vigorous physical activity in adults with prediabetes and type 2 diabetes. Buchowski M, editor. PLoS One. 2017;12: e0181053. doi:10.1371/journal.pone.0181053

19. Fishamn EI, Steeves JA, Zipunnikov V, Koster A, Berrigan D, Harris TA, et al. Association between Objectively Measured Physical Activity and Mortality in NHANES. Med Sci Sport Exerc. 2016;48: 1303–1311. doi:10.1249/MSS.0000000000000885

20. Chastin SFM, Palarea-Albaladejo J, Dontje ML, Skelton DA. Combined Effects of Time Spent in Physical Activity, Sedentary Behaviors and Sleep on Obesity and Cardio-Metabolic Health Markers: A Novel Compositional Data Analysis Approach. Devaney J, editor. PLoS One. 2015;10: e0139984. doi:10.1371/journal.pone.0139984

21. Carson V, Tremblay MS, Chaput J-P, Chastin SFM. Associations between sleep duration, sedentary time, physical activity, and health indicators among Canadian children and youth using compositional analyses 1. Appl Physiol Nutr Metab. 2016;41: S294–S302. doi:10.1139/apnm-2016-0026

22. Tremblay MS, Carson V, Chaput J-P, Connor Gorber S, Dinh T, Duggan M, et al. Canadian 24-Hour Movement Guidelines for Children and Youth: An Integration of Physical Activity, Sedentary Behaviour, and Sleep 1. Appl Physiol Nutr Metab. 2016;41: S311–S327. doi:10.1139/apnm-2016-0151

23. Paraschiv-Ionescu A, Buchser E, Aminian K. Unraveling dynamics of human physical activity patterns in chronic pain conditions. Sci Rep. 2013;3: 1–10. doi:10.1038/srep02019

24. Healy GN, Eakin EG, LaMontagne AD, Owen N, Winkler EAH, Wiesner G, et al. Reducing sitting time in office workers: Short-term efficacy of a multicomponent intervention. Prev Med (Baltim). 2013;57: 43–48. doi:10.1016/j.ypmed.2013.04.004

25. Healy GN, Dunstan DW, Salmon J, Cerin E, Shaw JE, Zimmet PZ, et al. Breaks in Sedentary Time: Beneficial associations with metabolic risk. Diabetes Care. 2008;31: 661–666. doi: 10.2337/dc07-2046

26. Healy GN, Matthews CE, Dunstan DW, Winkler EAH, Owen N. Sedentary time and cardio-metabolic biomarkers in US adults: NHANES 2003-06. Eur Heart J. 2011;32: 590–597. doi: 10.1093/eurheartj/ehq451

27. Chastin SFM, Ferriolli E, Stephens NA, Fearon KCH, Greig C. Relationship between sedentary behaviour, physical activity, muscle quality and body composition in healthy older adults. Age Ageing. 2012;41: 111–114. doi:10.1093/ageing/afr075

28. Lim ASP, Yu L, Costa MD, Buchman AS, Bennett DA, Leurgans SE, et al. Quantification of the Fragmentation of Rest-Activity Patterns in Elderly Individuals Using a State Transition Analysis. Sleep. 2011;34: 1569–1581. doi:10.5665/sleep.1400

29. Nakamura T, Kiyono K, Yoshiuchi K, Nakahara R, Struzik ZR, Yamamoto Y. Universal Scaling Law in Human Behavioral Organization. Phys Rev Lett. 2007;99: 138103. doi:10.1103/PhysRevLett.99.138103

30. Finkelstein M. Failure Rate Modelling for Reliability and Risk. London: Springer London; 2008. doi: 10.1007/978-1-84800-986-8

31. Johnson C, Paulose-Ram R, Ogden CL. National Health and Nutrition Examination Survey: Analytic guidelines, 1999-2010. Vital Heal Stat. 2013;2: 1–24.

32. Choi L, NLiu Z, Matthews CE, Buchowski MS. Validation of Accelerometer Wear and Nonwear Time Classification Algorithm. Med Sci Sport Exerc. 2011;43: 357–364. doi:10.1249/MSS.0b013e3181ed61a3

33. Troiano RP. Large-Scale Applications of Accelerometers. Med Sci Sport Exerc. 2007;39: 1501. doi:10.1097/mss.0b013e318150d42e

34. Tudor-Locke C, Camhi S, Troiano R. A Catalog of Rules, Variables, and Definitions Applied to Accelerometer Data in the National Health and Nutrition Examination Survey, 2003-2006. Prev Chronic Dis. 2012; doi:10.5888/pcd9.110332

35. Koster A, Caserotti P, Patel K V., Matthews CE, Berrigan D, Van Domelen DR, et al. Association of Sedentary Time with Mortality Independent of Moderate to Vigorous Physical Activity. Ruiz JR, editor. PLoS One. 2012;7: e37696. doi:10.1371/journal.pone.0037696

36. Troiano RP, Berrigan D, Dodd KW, Masse, Louise C, Tilert T, McDowell M. Physical Activity in the United States Measured by Accelerometer. Med Sci Sport Exerc. 2008;40: 181–188. doi: 10.1249/mss.0b013e31815a51b3

37. Lynch BM, Dunstan DW, Healy GN, Winkler E, Eakin E, Owen N. Objectively measured physical activity and sedentary time of breast cancer survivors, and associations with adiposity: findings from NHANES (2003-2006). Cancer Causes Control. 2010;21: 283–288. doi:10.1007/s10552-009-9460-6

38. Gastwirth JL. The Estimation of the Lorenz Curve and Gini Index. Rev Econ Stat. 1972;54: 306–316. doi:10.2307/1937992

39. Lim ASP, Yu L, Costa MD, Leurgans SE, Buchman AS, Bennett DA, et al. Increased Fragmentation of Rest-Activity Patterns Is Associated With a Characteristic Pattern of Cognitive Impairment in Older Individuals. Sleep. 2012;35: 633–640. doi:10.5665/sleep.1820

40. Lim ASP, Kowgier M, Yu L, Buchman AS, Bennett DA. Sleep Fragmentation and the Risk of Incident Alzheimer’s Disease and Cognitive Decline in Older Persons. Sleep. 2013;36: 1027–1032. doi:10.5665/sleep.2802

41. Lim ASP, Yu L, Kowgier M, Schneider JA, Buchman AS, Bennett DA. Modification of the Relationship of the Apolipoprotein E ε4 Allele to the Risk of Alzheimer Disease and Neurofibrillary Tangle Density by Sleep. JAMA Neurol. 2013;70: 1544. doi:10.1001/jamaneurol.2013.4215

42. Lee ET, Wang JW. Statistical Methods for Survival Data Analysi. volume 476. John Wiley & Sons, Inc.; 2003.

43. Miller R. Survival Analysis. Vol 66. John Wiley & Sons, Inc.; 2011.

44. Clauset A, Shalizi CR, Newman MEJ. Power-Law Distributions in Empirical Data. SIAM Rev. 2009;51: 661–703. doi:10.1137/070710111

45. Gillespie CS. Fitting Heavy Tailed Distributions: The poweRlaw Package. J Stat Softw. 2015;64. doi:10.18637/jss.v064.i02

46. Chastin SFM, Baker K, Jones D, Burn D, Granat MH, Rochester L. The pattern of habitual sedentary behavior is different in advanced Parkinson’s disease. Mov Disord. 2010;25: 2114–2120. doi:10.1002/mds.23146

47. Binder DA. Fitting Cox’s Proportional Hazards Models from Survey Data. Biometrika. 1992;79: 139–147. doi:10.2307/2337154

48. Lin D. On fitting Cox’s proportional hazards models to survey data. Biometrika. 2000;87: 37–47. doi:10.1093/biomet/87.1.37

49. Tsiatis AA. A Large Sample Study of Cox’s Regression Model. Ann Stat. 1981; 9: 93–108. doi:10.1214/aos/1176345335

50. Biswas A, Oh PI, Faulkner GE, Bajaj RR, Silver MA, Mitchell MS, et al. Sedentary Time and Its Association With Risk for Disease Incidence, Mortality, and Hospitalization in Adults. Ann Intern Med. 2015;162: 123–132. doi:10.7326/M14-1651

51. Wijndaele K, Brage S, Besson H, Khaw K-T, Sharp SJ, Luben R, et al. Television viewing time independently predicts all-cause and cardiovascular mortality: the EPIC Norfolk Study. Int J Epidemiol. 2011;40: 150–159. doi: 10.1093/ije/dyq105

52. Urbanek JK, Spira A, Di J, Leroux A, Crainiceanu C, Zipunnikov V. Epidemiology of Objectively Measured Bedtime and Chronotype in the US adolescents and adults: NHANES 2003-2006. 2017; Available: http://arxiv.org/abs/1706.05416

53. Bellettiere J, Winkler EAH, Chastin SFM, Kerr J, Owen N, Dunstan DW, et al. Associations of sitting accumulation patterns with cardio-metabolic risk biomarkers in Australian adults. Hu C, editor. PLoS One. 2017;12: e0180119. doi:10.1371/journal.pone.0180119

54. Chastin SFM, Egerton T, Leask C, Stamatakis E. Meta-analysis of the relationship between breaks in sedentary behavior and cardiometabolic health. Obesity. 2015;23: 1800–1810. doi:10.1002/oby.21180

